# USP7 inactivation suppresses APC-mutant intestinal hyperproliferation and tumor development

**DOI:** 10.1101/2022.09.22.508986

**Authors:** Laura Novellasdemunt, Anna Kucharska, Anna Baulies, Georgios Vlachogiannis, Dimitra Repana, Andrew Rowan, A Suárez-Bonnet, Francesca Ciccarelli, Nicola Valeri, Vivian S. W. Li

**Author notes:** Correspondence should be addressed to Vivian Li.

## Abstract

Truncating mutation of the tumor suppressor gene adenomatous polyposis coli (APC) is the hallmark of colorectal cancer (CRC), resulting in constitutive WNT activation. Despite decades of research, targeting WNT signaling in cancer remains challenging due to its essential role in normal stem cell maintenance. We have previously shown that the deubiquitinating enzyme USP7 is a tumor-specific WNT activator in APC-truncated cells by deubiquitinating and stabilizing β-catenin, but its role in gut tumorigenesis is unknown. Here we show *in vivo* that deletion of Usp7 in Apc-truncated mice inhibits crypt hyperproliferation and intestinal tumor development. Importantly, intestine-specific Usp7 mutation does not yield any phenotype in wildtype animals, indicating that its loss is well tolerated. Unexpectedly, prolonged deletion of Usp7 in Apc+/− intestine induces varying degrees of colitis. Treatment with a USP7 inhibitor suppresses growth of patient-derived cancer organoids *in vitro* and of xenografts carrying APC truncations. We propose that USP7 inhibition may be efficacious for tumor-specific therapy of sporadic APC-mutated CRC, while patients with germline APC mutations should not receive such treatment.

**Highlights:**

- Usp7 deletion in Apc-truncated mice reduces intestinal tumor development.
- Intestine-specific Usp7 mutation mutation has no phenotype in wildtype animals.
- Treatment with Usp7 inhibitor suppresses growth of patient-derived cancer organoids carrying Apc truncations *in vitro* and of xenografts.

## INTRODUCTION

Colorectal cancer (CRC) is the third most commonly diagnosed cancer and the second cause of cancer-related death worldwide (1). CRC can be classified into two major categories: the microsatellite instable (MSI) subtype, characterized by defective mismatch repair machinery and hypermutations, and the non-hypermutated microsatellite stable (MSS) subtype. The majority of CRCs (∼85%) are MSS, characterized by sequential acquisition of genetic alterations including *APC, KRAS, TP53* and *SMAD4* (2). Adenomatous polyposis coli (*APC*) mutation is the first step for tumor initiation in MSS CRC, leading to hyperactivation of the WNT signaling pathway (3–5). The canonical WNT/β-catenin pathway regulates the levels of the Wnt effector protein β-catenin through phosphorylation and ubiquitination-mediated degradation in the cytoplasmic β-catenin destruction complex. The latter consists of APC, AXIN, glycogen synthase kinase 3 (GSK3), and casein kinase 1 (CK1). In the absence of WNT ligands, β-catenin is sequentially phosphorylated by CK1 and GSK3 followed by recruitment of the E3 ubiquitin ligase β-TrCP to the destruction complex for ubiquitination and subsequent proteasomal degradation (6–8). Engagement of WNT ligands to the receptors inhibits the β-catenin destruction complex, leading to accumulation of β-catenin in the cytoplasm and nucleus for transcriptional activation of the WNT target genes (9).

Previous studies showed that deletion of *APC* in mice leads to crypt hyperproliferation and adenoma formation in the intestine through constitutive activation of WNT signaling and β-catenin/TCF transcription of target genes (10,11). In human, somatic and germline mutations of *APC* were discovered in CRC patients in 1991 (12–15), where the majority of somatic *APC* mutations occur in the “mutation cluster region” (MCR) between codons 1286 and 1513 (2, 16). Region-specific *APC* mutations have been associated with differential WNT/β-catenin transcriptional activity and tumor susceptibility (17). We have previously shown that *APC* truncating mutations activate WNT signaling through abrogation of β-catenin ubiquitination within the destruction complex in CRCs (9). Despite decades of research, safe drugs that correct *APC* loss in (colon) cancer remain elusive.

A number of inhibitors targeting upstream components of the WNT pathway (e.g. the WNT receptors Frizzled (18–21) and the WNT ligand secretory machinery: porcupine (22)) are currently in clinical trials. However, these inhibitors are not effective in *APC*-mutated CRC where constitutive WNT activation is independent of ligand-receptor activation. Tankyrase inhibitors and β-catenin/CPB antagonists acting in the cytoplasm and nucleus respectively have been previously proposed to suppress WNT signaling in *APC*-mutated CRC cells (23–28). However, these inhibitors exhibit high on-target toxicity due to the pivotal role of WNT signaling in other healthy tissues (29). This poses safety concerns for the use of the WNT inhibitors in the clinic. For many years, the WNT pathway has been considered as undruggable.

We have recently found that a deubiquitinating enzyme USP7 is responsible for WNT activation specifically in *APC*-mutated CRC (30). We showed that USP7 is recruited to the APC-truncated destruction complex for β-catenin deubiquitination and stabilization. Deletion of USP7 in cancer cell lines and organoids inhibits WNT activation by restoring β-catenin ubiquitination specifically in *APC*-mutated but not wild-type (WT) cells, suggesting that USP7 may represent a tumor-specific drug target. Contrary to our finding, a more recent study showed that USP7 is a potent negative regulator of WNT/β-catenin signaling by deubiquitinating and stabilizing AXIN (31), casting doubt on the use of USP7 as WNT inhibitor for *APC*-mutated CRC patients. Importantly, the functional role of USP7 in CRC has not yet been explored *in vivo*.

Here, we investigate the role of USP7 in tumor development and intestinal homeostasis *in vivo*. Comprehensive analyses of three different *Apc*-deletion mouse models show that loss of Usp7 significantly reduces crypt hyperproliferation, WNT activation and tumor development in *Apc*-deficient intestine. Consistent with our previous observation in cancer cell lines and organoids, deletion of *Usp7* in the Apc WT animals does not perturb intestinal homeostasis. We further validate the findings in human CRC, where treatment with a USP7 inhibitor suppresses growth in both patient-derived organoids (PDOs) and xenografts carrying APC truncating mutations, supporting the notion that USP7 can be used as a tumor-specific drug target for *APC*-mutated CRCs by suppressing WNT signaling.

## RESULTS

### Usp7 depletion suppresses intestinal crypt hyperproliferation and WNT activation mediated by homozygous loss of APC

To explore the therapeutic role of USP7 in CRC, we first examined the effect of *Usp7* deletion in an acute WNT activation model mediated by homozygous loss of Apc. Villin^CreERT2^;Apc^fl/fl^ (Apc^fl/fl^) mice (10) were crossed with Villin^CreERT2^;Usp7^fl/fl^ mice (32) to obtain intestinal-specific conditional deletion of *Apc* and *Usp7*. Homozygous loss of Apc caused hyperproliferation of the intestinal crypts within 6 days after tamoxifen induction (Fig. 1a-b, 1d-e). This was accompanied by increased expression of the stem cell marker *Lgr5* (33) (Fig. 1g-h) and WNT targets Cyclin D1 and Sox9 (Fig. 1j-k, 1m-n) compared to WT control (Villin^CreERT2^) intestine, and loss of differentiation markers as indicated by Periodic Acid (PAS) staining (goblet cells) (Fig. 1 p-q) and Keratin 20 (Fig. 1s-t). Strikingly, deletion of Usp7 in Apc^fl/fl^ intestine (Apc^fl/fl^ Usp7^fl/fl^) resulted in a marked suppression of crypt hyperproliferation (Fig. 1c, 1f, and Fig.S1a) and a significant reduction in the size of the crypts (Fig. S1b) and the length of the crypt-villus axis (Fig. S1c) throughout the gut. In addition, the stem cell marker Lgr5 (Fig. 1i) and WNT targets Cyclin D1 and Sox9 (Fig. 1l, 1o) were consistently suppressed in the Apc^fl/fl^ Usp7^fl/fl^ intestine, while differentiation was partially restored (Fig. 1r, 1u and S1d).

**Figure 1.**
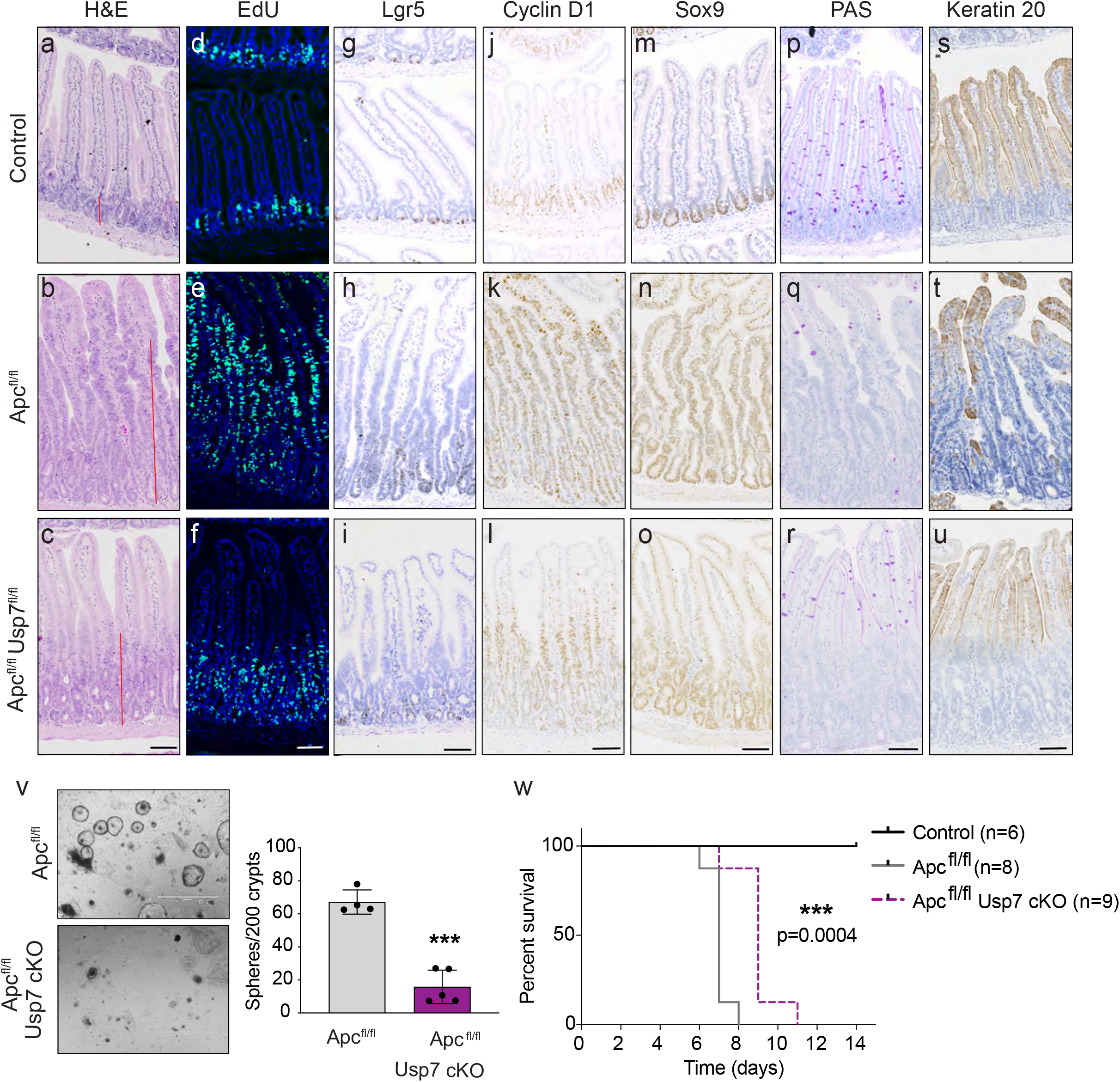
Loss of Usp7 ameliorates acute WNT activation phenotype. (**a-u**) Histology and immunostaining using the indicated markers (n=3 per group). Representative images of H&E staining (a-c), Edu (d-f), Lgr5 RNAscope (g-i), Cyclin D1 (j-l), Sox9 (m-o), PAS (p-r) and Keratin 20 staining (s-u) from Villin^CreERT2^ (Control), Villin^CreERT2^;Apc^fl/fl^ (Apc^fl/fl^) and Villin^CreERT2^;Apc^fl/fl^;Usp7^fl/fl^ (Apc^fl/fl^ Usp7^fl/fl^) intestine at 6 dpi. Scale bars, 50 μm. (**v**) Representative images of colony formation assay of organoids derived from Apc^fl/fl^ and Apc^fl/fl^ Usp7^fl/fl^ intestines at day 6 dpi. Scale bar, 1000 μm. Right, quantitation of the number of spheres form after seeding 200 crypts. Data are mean ± standard error. At least n=4 per group. P-values were determined using the unpaired two-sided t-test (***P<0.001). (**w**) Kaplan-Meier survival analysis of Control, Apc^fl/fl^ and Apc^fl/fl^ Usp7^fl/fl^ mice. P-values were determined using the Mantel-Cox test.

To validate the WNT inhibition and growth suppression phenotype, we further examined the intestinal organoids derived from Apc^fl/fl^ and Apc^fl/fl^ Usp7^fl/fl^ animals. Consistent with the *in vivo* data, Usp7 depletion strongly suppressed the organoid formation efficiency in the Apc-deficient background (Fig. 1v). Quantitative RT-PCR analysis confirmed the loss of Usp7 transcripts in organoids (Fig. S1e), where WNT target genes (*Axin2* and *Sox9*) and stem cell markers (*Olfm4* and *Ascl2*) were significantly downregulated in the Apc^fl/fl^ Usp7^fl/fl^ organoids (Fig. S1f). Together, the results indicate that Usp7 deletion in the intestine suppresses WNT signaling and crypt hyperproliferation mediated by *Apc* mutation.

Interestingly, deletion of Usp7 prolonged the survival of the Apc^fl/fl^ mice from an average of 7 days to 9 days but not beyond (Fig. 1w). This is likely due to the highly aggressive phenotype of the acute WNT activation model driven by homozygous loss of Apc, where the entire gut epithelium is acutely transformed into adenomatous tissue.

### Usp7 is dispensable for intestinal homeostasis in wild-type mice

Our previous data using CRC cell lines and organoids showed that USP7 is essential for pathological WNT activation in *APC* mutant cells but not physiological WNT signaling in normal cells, suggesting that USP7 can be used as tumor-specific drug target (30). However, an independent study showed that USP7 functions as a negative regulator of WNT, where inhibition of USP7 increases WNT signaling in normal cells (31). To clarify the role of USP7 in normal intestine, we generated Villin^CreERT2^;Usp7^fl/fl^ (Usp7 cKO) mice to induce Usp7 deletion specifically in the intestine upon tamoxifen administration. Usp7-depleted intestine did not show any noticeable histological differences, including crypt length (Fig. 2a-b and 2k) or cell proliferation (Fig. 2c-d and 2l) 17 days post-induction (dpi). Similarly, expression of the stem cell marker Olfm4 (Fig. 2e-f and 2m), WNT target Cyclin D1 (Fig. 2g-h and 2n) and goblet cell differentiation (Fig. 2i-j and 2o) were unaltered. We further examined if p53 is activated upon Usp7 deletion since it is also a proposed target of USP7 (34, 35). Immunostaining of the Usp7 cKO intestine did not show any induction of p21 expression (Fig. S2a), indicating that Usp7 loss does not activate p53-mediated growth arrest in the intestine.

**Figure 2.**
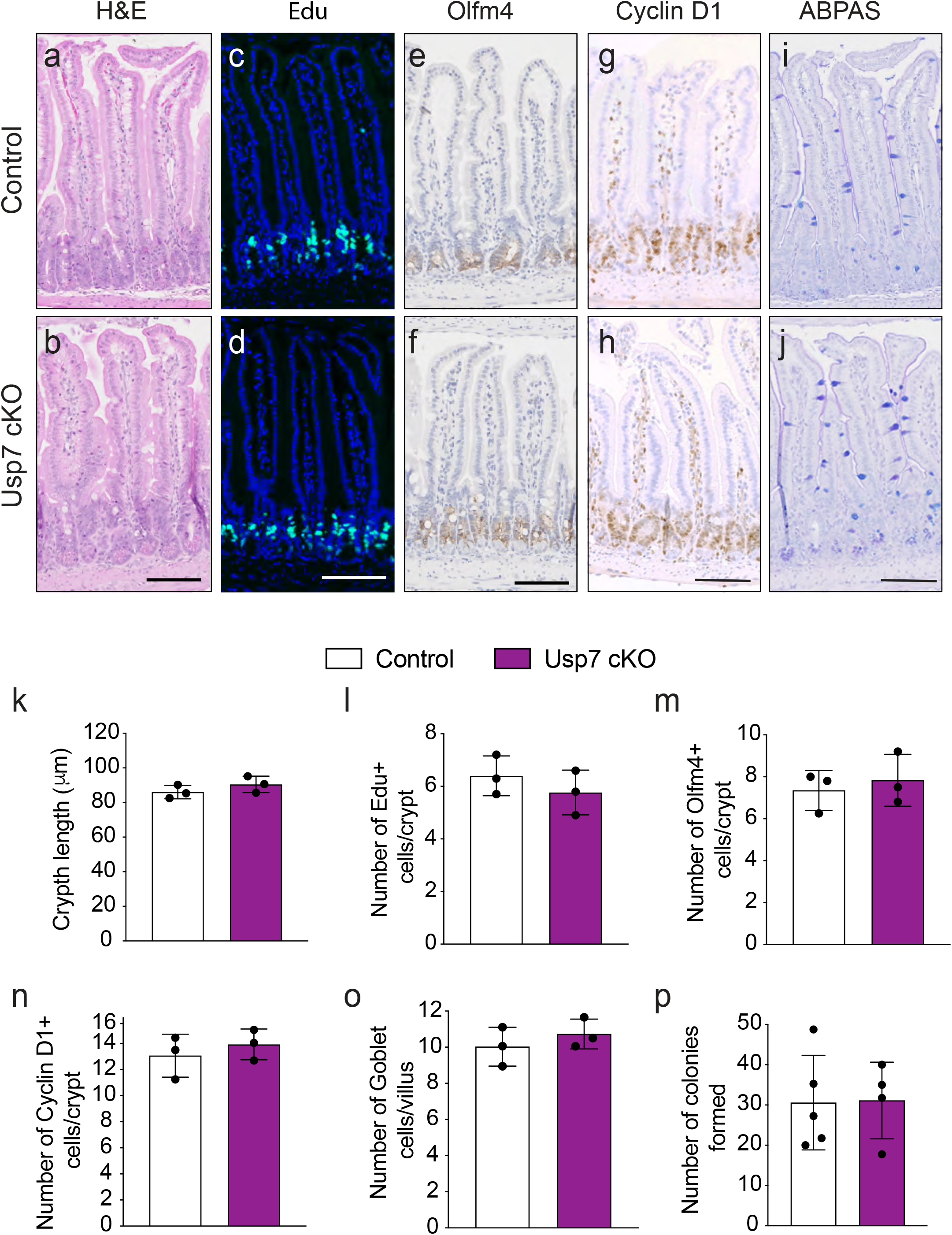
Usp7 inhibition in wild-type mice does not perturb intestinal homeostasis. (**a-j**) Histology and immunostaining using the indicated markers. n=3 per group. Representative images of H&E staining (a-b), Edu (c-d), Olfm4 (e-f), Cyclin D1 (g-h) and AB-PAS (i-j) from Villin^CreERT2^ (Control) and Villin^CreERT2^;Usp7^fl/fl^ (Usp7 cKO) intestine at 17 dpi. Scale bars, 50 μm. (**k**) Quantitation of crypt length (μm) from (a-b). Each dot represents the average of at least 20 crypts per animal. Data are mean ± standard error. n=3 per group. (**l-o**) Quantitation of Edu positive cells (l) from (c-d), Olfm4 (m) from (e-f), Cyclin D1 (n) from (g-h) and AB-PAS staining (indicative of Goblet cells) (o) from (i-j). Each dot represents the average of at least 20 crypts per animal (except for Edu that is 10 crypts per animal). Data are mean ± standard error. n= 3 per group. P-values were determined using the unpaired two-sided t-test (*P<0.05). (**p**) Quantitation of the organoid formation assay in organoids derived from control and Usp7 cKO mice. Data are mean ± standard error. Control, n=5; Usp7 cKO, n=3.

Next, we established organoids from the control and Usp7 cKO intestine, which showed no differences in morphology (Fig. S2b) or organoid formation efficiency (Fig. 2p), suggesting that Usp7 deficiency does not affect intestinal stem cell proliferation. Indeed, unlike in the Apc^fl/fl^ background, loss of Usp7 in WT organoids did not alter the expression of stem cell, WNT target genes or differentiation markers (Fig. S2c-f). The results demonstrate that deletion of Usp7 alone does not alter WNT signaling nor tissue homeostasis in normal WT intestine.

To further study the long-term effect of Usp7 deletion, Usp7 cKO animals were aged for 1.5 years followed by tissue analysis. Consistent with the short-term deletion data, long-term loss of Usp7 did not cause any significant changes in tissue morphology (Fig. S2g-h), proliferation (Fig. S2i-j), and expression of stem cell markers (Fig. S2k-l) and WNT target genes (Fig. S2m-p). Quantitative RT-PCR analysis of the intestinal crypts isolated from these mice further confirmed that the stem cell markers and WNT target gene expression remained unchanged after long-term deletion of Usp7 (Fig. S2q-s). Our data support the notion that loss of Usp7 alone in WT mice does not impact WNT signaling or intestinal stem cell proliferation.

### Usp7 depletion suppresses intestinal tumor development in Apc^min^ and Apc^fl/+^ mice

Next, we investigated the role of Usp7 in intestinal tumorigenesis. The Apc^min^ mouse strain, modeling human Familial Adenomatous Polyposis (FAP), was used as a chronic spontaneous model for intestinal tumor development (36). Comparison of Apc^min^ and Apc^min^ Usp7 cKO mice showed that Usp7 deletion significantly reduced tumor numbers at 4 months of age (Fig. 3a-b). Histological analysis revealed that both low-grade and high-grade dysplasias were present in all Apc^min^ control animals, whereas the majority of the adenomas in the Apc^min^ Usp7 cKO mice were low-grade dysplasias (Fig. 3c). The results indicate that Usp7 deficiency inhibits tumor progression in Apc^min^ mice. Loss of Usp7 was confirmed by quantitative RT-PCR analysis of the isolated crypts of the animals (Fig. S3a). Immunostaining of cleaved-Caspase3 did not show any difference between Apc^min^ Control and Apc^min^ Usp7 cKO tumors, indicating that the tumor reduction upon Usp7 deletion was not caused by increased apoptosis (Fig. S3b). Quantitative RT-PCR analysis of the isolated tumors showed that expression of the WNT target genes and stem cell markers *Lgr5, Ascl2* and *Axin2* were suppressed in the Usp7 cKO tumors (Fig. 3d), suggesting that Usp7 depletion inhibits WNT signaling in the Apc-deficient intestinal tumors. Interestingly, histological analysis of the established tumors in both Apc^min^ and Apc^min^ Usp7 cKO mice showed reduced proliferation as assessed by Edu staining (Fig. S3c) and decreased stem cell marker Sox9 expression (Fig. S3d). Surprisingly, unlike our observations at earlier time points, we observed no difference in the survival of Apc^min^ Usp7 cKO mice or their overall tumor burden compared to Apc^min^ control (Fig. 3e, Fig. S3e-f). It is worth noting that the Apc^min^ Usp7 cKO tumors showed incomplete depletion of Usp7 (Fig. 3d), suggesting some degree of escape phenotype in these tumours. Importantly, histological examination of the intestinal tissues revealed that all Apc^min^ Usp7 cKO animals displayed various degree of colitis pathology (Fig. 3f-g) as well as enteritis (Fig. 3h), which may have worsened the health of the animals. The data suggest that, although loss of Usp7 does not alter tissue homeostasis in WT intestine, there might be a deleterious effect when Usp7 is inhibited in intestinal cells carrying only one copy of WT *Apc* gene (as in the Apc^min^ mouse model) due to haplo-insufficiency. We speculate that the colitis and enteritis observed in the Apc^min^ Usp7 cKO intestine may promote inflammation-associated CRC development over time (37), which may compensate the tumor suppressive effect of Usp7 loss in these animals.

**Figure 3.**
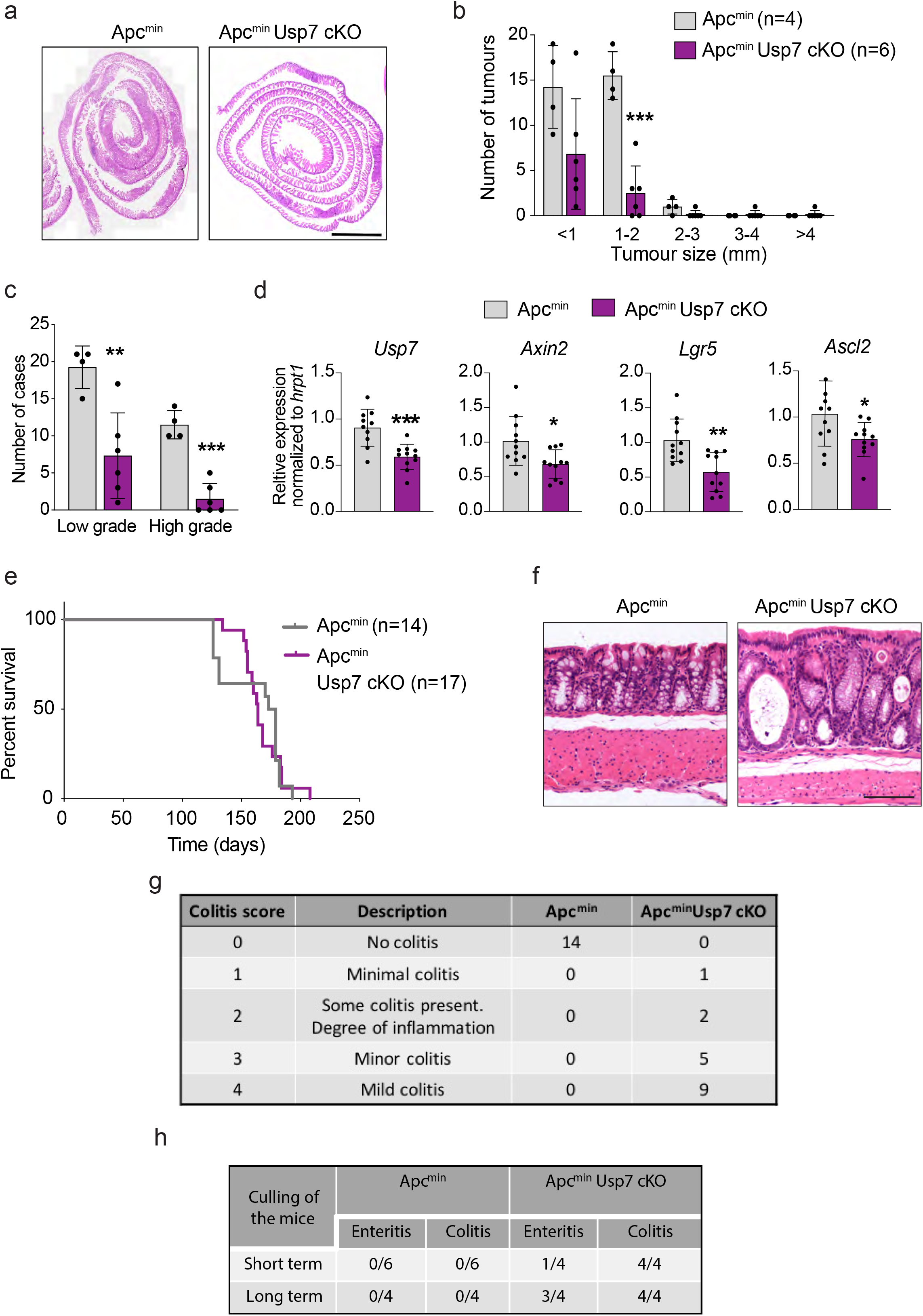
Loss of Usp7 reduces tumor number in Apc^min^ mice. (**a**) Representative H&E stainings of the small intestines of the indicated genotypes. Scale bar, 2000 μm. (**b**) Total number of adenomas 120 days after induced Usp7 loss. Data are mean ± standard error. Apc^min^ control n=4 and Apc^min^ Usp7 cKO n=6. (**c**) Quantitation of the grades of the adenomas that developed in the indicated mice. Data are mean ± standard error. Apc^min^ control n=4 and Apc^min^ Usp7 cKO n=6. (**d**) mRNA expression of the indicated genes was analyzed by qRT-PCR in tumors isolated from mice with the indicated genotypes (n=3 per condition). Data are presented as fold change normalized to *hrpt1* control. Error bars represent ± standard error. (**e**) Kaplan-Meier survival analysis of Control Apc^min^ and Apc^min^ Usp7 cKO mice. P-value was determined using the Mantel-Cox test. (**f**) Representative H&E staining from Apc^min^ control and Apc^min^ Usp7 cKO mice showing colitis present in the Apc^min^ Usp7 cKO intestine. Scale bar, 100 μm. (**g**) Table summarizing the number of mice that developed colitis in the indicated genotypes. (**h**) Table depicting the number of mice that developed colitis and enteritis in the indicated genotypes and time. Short term refers at around day 90 dpi and long term refers at around day 200 dpi. All mice showing enteritis where of grade 2-3 (grading description in methods).

To validate the hypothesis of Apc haplo-insufficiency in Usp7 KO background, we generated another Apc heterozygous model using Villin^CreERT2^;Apc^fl/+^ (Apc^het^) animals and further crossed to Usp7 cKO mice. Tamoxifen was injected to the Apc^het^ and Apc^het^ Usp7 cKO animals, and intestinal tissues were harvested at around 90 dpi for histological analysis. Remarkably, adenomas were observed in all 6 Apc^het^ mice, while none of the Apc^het^ Usp7 cKO mice developed any tumor (Fig. 4a-c). This is consistent with our earlier observation that loss of Usp7 inhibits Apc-deficient tumor development. Quantitative RT-PCR analysis confirmed Usp7 depletion as well as reduced expression of stem cell markers (Lgr5 and Olfm4) and WNT target gene (Sox9) in the Apc^het^ Usp7 cKO intestine (Fig. 4d). Immunostaining showed aberrant expression of Sox9, Cyclin D1 and Edu in the Apc^het^ tumors (Fig. S4a,c,e), whilst expression of the WNT targets and proliferation markers remained restricted to the untransformed crypts of the Apc^het^ Usp7 cKO intestine (Fig. S4b,d,f). Similar to the Apc^min^ model, we observed no difference in the survival and tumor burden of the Apc^het^ Usp7 cKO mice at later time points (Fig. 4e, Fig. S4g-h). We also observed colitis pathology in all of the Apc^het^ Usp7 cKO mice (n=11), while only 2 out of 13 Apc^het^ control animals showed minor degree of inflammation (Fig. 4f-g). In addition, enteritis was also observed in 2 out of 5 Apc^het^ Usp7 cKO animals culled at short term (around day 90), and was more abundant (4 out of 5 animals) at long term (day 200) (Figure 4h). The data support the notion of inflammation-associated tumorigenesis in the Apc^het^ Usp7 mice.

**Figure 4.**
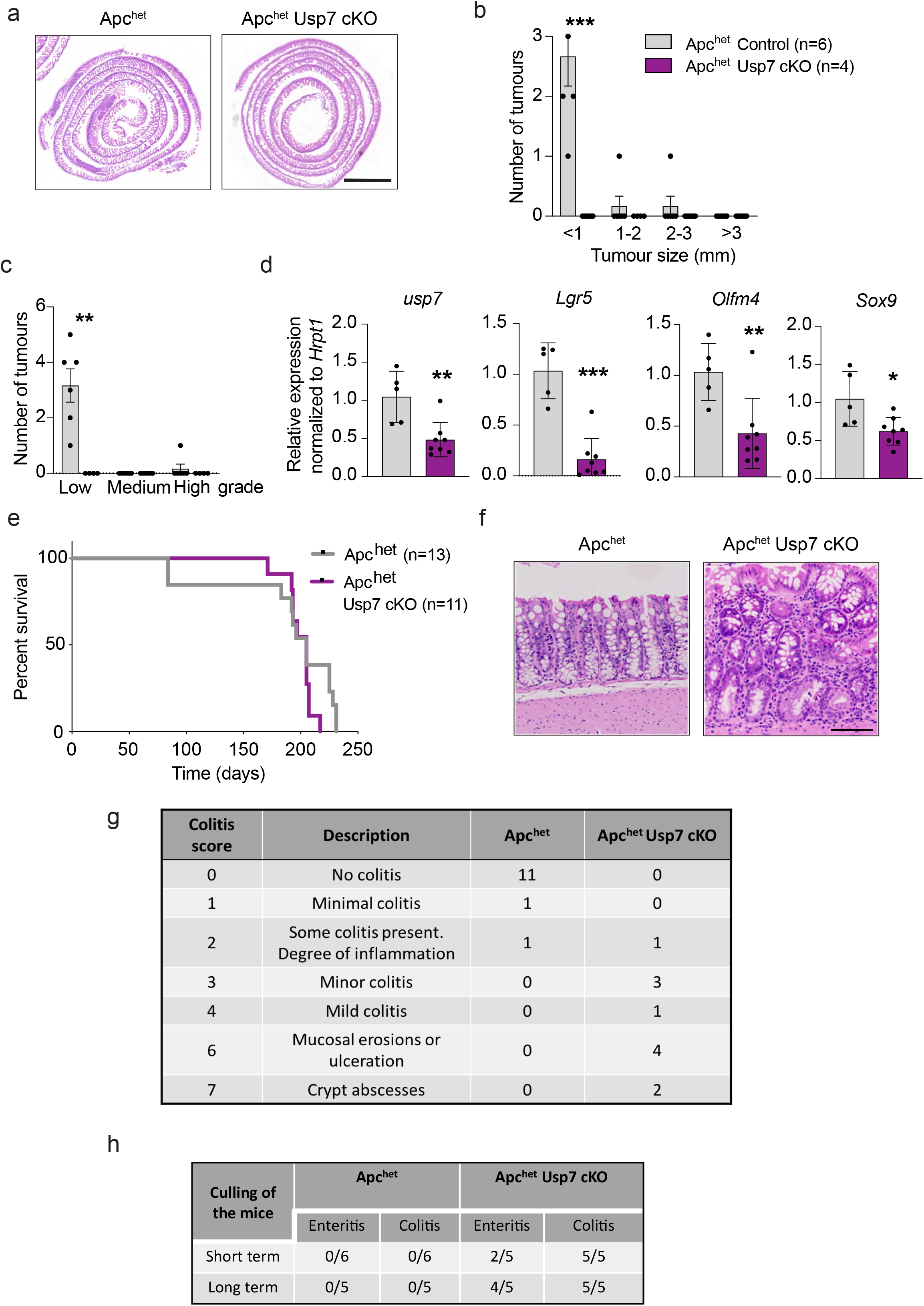
Usp7 deletion inhibits tumor growth in Apc^het^ mice. (**a**) Representative H&E stainings of the small intestines of the indicated genotypes. Scale bar, 2000 μm. (**b**) Total number of adenomas 84 days after induced Usp7 loss. Data are mean ± standard error. Apc^het^ control n=6 and Apc^het^ Usp7 cKO n=4. Error bars represent ± standard error. P-values were determined using the unpaired two-sided t-test (***P<0.001). (**c**) Quantitation of the grades of the adenomas that developed in the indicated mice. Data are mean ± standard error. Apc^het^ control n=6 and Apc^het^ Usp7 cKO n=4. Error bars represent ± standard error. P-values were determined using the unpaired two-sided t-test (**P<0.01). (**d**) mRNA expression of the indicated genes was analyzed by qRT-PCR in tumors isolated from mice with the indicated genotypes (n=3 per condition). Data are presented as fold change normalized to *hrpt1* control. Error bars represent ± standard error. (**e**) Kaplan-Meier survival analysis of Control, Apc^het^ and Apc^het^ Usp7 cKO mice. P values were determined using the Mantel-Cox test. (**f**) Representative H&E staining from Apc^het^ control and Apc^het^ Usp7 cKO mice showing colitis present in the Apc^het^ Usp7 cKO mice. Scale bar, 100 μm. (**g**) Table summarizing the number of mice that developed colitis in the indicated genotypes. (**h**) Table summarizing the number of mice that developed colitis and enteritis in the indicated genotypes and time. Short term refers at around day 90 dpi and long term refers at around day 200 dpi. All mice showing enteritis where of grade 2-3 (grading description in methods).

Altogether, these results indicate that inhibition of Usp7 suppresses intestinal tumor development and progression mediated by *Apc* truncating mutations. Importantly, our data show that Usp7 depletion is well-tolerated in normal intestine but may have an adverse effect on Apc heterozygous (Apc+/−) intestinal cells due to haplo-deficiency. This implies that USP7 can potentially be an effective therapeutic target for sporadic *APC*-mutated CRCs, whilst FAP patients carrying germline *APC* mutations may be susceptible to colitis when treated with USP7 inhibitors.

### USP7 inhibition suppresses WNT signaling and growth of human colorectal cancer patient-derived organoids carrying *APC* truncating mutations

To validate our findings in human CRC, we further tested the therapeutic potential of USP7 by treating PDOs with the USP7 inhibitor P22077 that has been previously used for *in vivo* studies (30, 38). Human PDOs were established from 5 CRC patients carrying different *APC* mutations (Fig. 5a). PDOs 1-3 harbor different truncating *APC* mutations, all lacking the β-catenin inhibitory domain (CID) critical for pathological WNT activation and tumor transformation via USP7 binding (30). PDO4 carries a splice variant mutation at the −8 position upstream of exon 8 of the *APC* gene, leading to early frameshift and premature truncation (39). PDO5 has 2 missense mutations of *APC* after the CID domain towards the 3’ end that are predicted to yield full-length (FL), non-pathological APC protein. Microsatellite repeat analysis showed that PDO5 should be classified as an MSI CRC with WT APC and low WNT activity.

**Figure 5.**
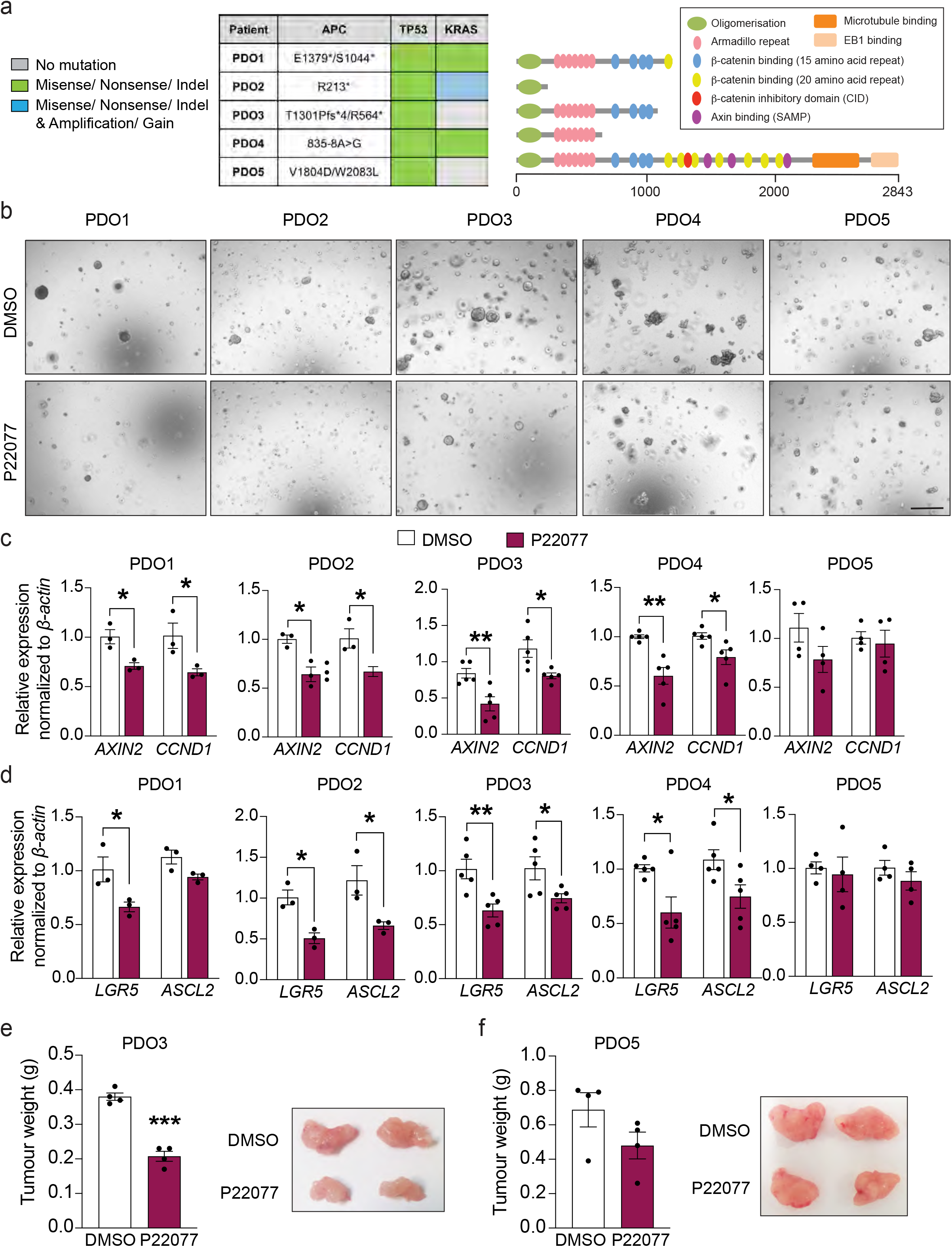
USP7 inhibition suppresses WNT signaling and tumor growth in human CRC patient-derived organoids carrying *APC* truncating mutations. **a**) Left: Mutation status of the key driver genes in PDOs 1-5. *indicates truncated protein. Right: Schematic representation of the APC protein products with different domains depicted. Numbers indicate codon numbers. (**b**) Representative images of colony formation assay of organoids treated with P22077 or vehicle DMSO. Scale bar, 500 μm. (**c**-**d**) mRNA expression of the indicated genes was analyzed by qRT-PCR in the organoids from (b). Data are presented as fold change normalized to *β-actin* control (at least n=4 per condition). (**e**-**f**) Tumor weights of PDO3 (e) and PDO5 (f) between DMSO control and P22077 treatment group (30 mg/kg) was measured at the end of treatment (20 days) (n=4 per condition). Representative images of tumors derived from PDO3 (e) and PDO5 (f) at the end of treatment. Error bars represent ± standard error. P-values were determined using the unpaired two-sided t-test (*P<0.05; **P<0.01; ***P<0.001).

Treatment of the PDOs with P22077 resulted in a growth reduction in the APC-truncated PDOs 1-4 but not the APC-FL PDO5 (Fig. 5b and Fig. S5a). Consistently, organoid formation efficiency was also reduced in PDOs 1-4 (Fig. S5b). Quantitative RT-PCR analysis confirmed that the WNT target genes (*AXIN2* and *CCND1*) and stem cell markers (*LGR5* and *ASCL2*) were significantly reduced in PDOs 1-4 upon USP7 inhibition, while PDO5 was unaffected by the treatment (Fig. 5c-d). These results are in concordance with the *in vivo* mouse data that USP7 inhibition suppresses hyperproliferation and tumor development specifically in APC-deficient intestine.

Next, we explored if USP7 inhibition could be used as an adjuvant therapy to fluorouracil (5-FU), a common chemotherapy given to CRC patients in the clinic. Cell Titer Glo viability analysis showed that the 5-FU-mediated growth suppression efficiency varied amongst the 5 PDOs, where PDO3 and PDO4 responded better than PDOs 1, 2 and 5 (Fig. S5c-d). Combination treatment of 5-FU and P22077 showed additive growth suppression as compared to 5-FU alone in PDOs 2-4 (Fig. S5c). In particular, the organoid growth of PDO3 and PDO4 was nearly completely abolished in the combined treatment (Fig. S5c-d). On the other hand, P22077 treatment did not further increase the 5-FU-mediated growth suppression in PDO5. Similar results were observed when analyzing organoid formation capacity (Fig. S5e). These data suggest that USP7 inhibition may be used in combination with current chemotherapeutic drugs such as 5-FU in *APC*-mutated CRCs.

Finally, we tested the therapeutic potential of USP7 inhibitor *in vivo* by transplanting the PDOs subcutaneously into immunodeficient mice. To validate the *in vitro* treatment data, the good (PDO3) and poor (PDO5) responders of the USP7 inhibitor were selected for the xenograft experiment. USP7 inhibitor P22077 or vehicle control (DMSO) were administered daily by intraperitoneal injections (30). Treatment by P22077 of the PDO3-xenografts significantly suppressed tumor growth compared to vehicle treatment (Fig. 5e). In contrast, PDO5-xenografts did not show significant effect on tumor development upon P22077 treatment (Fig. 5f). Similar to our previous observation (30), treatment of the USP7 inhibitor *in vivo* did not show any detectable health problems or weight loss (Fig. S6a). In addition, histological analysis of the P22077-treated PDO3-tumors showed a decreased expression of the WNT target gene CCND1 (Fig. S6b). Quantitative RT-PCR analysis of PDO3 further confirmed downregulation of *CCND1* and *AXIN2* upon P22077 treatment (Fig. S6c). On the other hand, expression of CCND1 and AXIN2 was unchanged in the P22077-treated PDO5-tumors (Fig. S6d-e). Of note, cleaved caspase3 expression was unaltered in both PDO-xenografts upon treatment, suggesting that P22077-mediated tumor suppression is not caused by increased apoptosis (Fig. S6f). Altogether, the data support the notion that USP7 inhibition can be used for treatment of *APC*-mutated CRC.

## DISCUSSION

The major genetic events that cause CRC have been extensively characterized over the past decades. In particular, loss-of-function mutations of *APC* represent the key driver mutation for over 80% of CRC. It has been previously shown that CRCs carrying *KRAS* and *p53* mutations remain strictly dependent on APC loss (40), indicating that the WNT signaling pathway is an important therapeutic target for treatment of CRC. However, despite decades of research, no approved drugs targeting WNT signaling in *APC*-mutated cancer exist. One of the major challenges is the crucial role of the WNT pathway in adult tissue homeostasis, making it difficult to develop safe and effective WNT inhibitors for treating CRC without on-target toxicity (5, 41). There is an urgent unmet need to develop a new generation of WNT inhibitors that exhibit tumor-specificity.

We have recently identified USP7 as a tumor-specific target in CRC carrying APC truncating mutations (30). We showed that APC truncations lacking the CID make β-catenin vulnerable to USP7 deubiquitination: depletion of USP7 in *APC* mutant CRC cell lines and mouse organoids inhibits WNT and suppresses cell growth. Importantly, we demonstrated that the USP7 is dispensable for WNT activity and cell viability in APC-WT cells, highlighting its therapeutic potential for CRC treatment.

Contrasting with our findings, it has been recently reported that USP7 functions as a potent negative WNT regulator by deubiquitinating AXIN in normal HEK293T cells, mesenchymal stem cells C3H10T1/2 and bone marrow-derived stroma cells ST2 (31). This raises uncertainty about the roles of USP7 in WNT regulation and the safe use of USP7 inhibitors for treating CRC. To address these concerns, we have generated a number of transgenic mouse strains to test the functional role of Usp7 in adult intestinal homeostasis and tumorigenesis *in vivo*. Both short-term and long-term intestine-specific loss of Usp7 reveals no major phenotype in WT animals, supporting the notion that Usp7 is dispensable for WNT signaling and tissue homeostasis in the normal WT intestine. In contrast, Usp7 depletion in Apc^fl/fl^ mice ameliorates the acute crypt proliferation and survival, with reduced expression of WNT target genes and stem cell markers accompanied by increased differentiation. The data is consistent with a previous finding whereby restoring APC function promotes tissue differentiation and tumor regression (40).

We further investigated the role of Usp7 in CRC using two independent mouse intestinal tumor models (Apc^min^ and Apc^het^) that mimic human FAP patients with germline *APC* mutations predisposed to CRC development. Loss of Usp7 significantly reduces tumor numbers and tumor grade in both models, indicating that Apc-deficient tumor development and progression is Usp7-dependent. This is further validated in human PDOs where treatment of USP7 inhibitor suppresses the growth of APC-truncated but not APC-FL CRCs, supporting the idea of using USP7 inhibitors for CRC treatment. Nevertheless, the survival of both mouse Apc+/−tumor models was not improved despite the complete absence of tumor development in Apc^het^ Usp7 cKO animals at 90 dpi. To our surprise, we observed intestinal tumor development at later time points in both Apc^min^ Usp7 cKO and Apc^het^ Usp7 cKO animals. Histological analysis showed colitis and enteritis inflammation in both Apc^min^ Usp7 cKO and Apc^het^ Usp7 cKO mice at approximately 3-4 months of age, which was not detected in the WT Usp7 cKO animals even after 1.5 years deletion. The results suggest that Usp7 inhibition is well tolerated in APC WT normal cells but may cause toxicity in APC heterozygous cells. Conceivably, heterozygous loss of APC may partially expose β-catenin to USP7, resulting in a moderate dependency on USP7 in those cells. We propose that USP7 inhibitors can potentially be effective in treating sporadic *APC*-mutated CRCs lacking the CID domain (30), while FAP patients with germline *APC* mutations should not receive such treatment due to the potential risk of developing colitis and the associated tumorigenesis. Further studies will be needed to understand the colitis development in the Apc^min^/Apc^het^ Usp7 cKO mice.

USP7 has been previously shown to regulate p53-dependent apoptosis by controlling the levels of p53 and Mdm2 (34, 35). Our previous study showed that USP7-mediated WNT activation is p53-independent (30). Indeed, we did not observe any transactivation of p21 protein expression in the Usp7 cKO intestine. Importantly, p53 is mutated in all human PDOs used in this study regardless of their response to the USP7 inhibitor treatment, which is consistent with our previous observation that the WNT-regulatory role of USP7 is p53-independent. Our PDO treatment data further suggest that USP7 inhibition can be used either alone or in combination with chemotherapeutic agents already being utilized clinically in treating CRC. It will be interesting to explore if p53 plays a role in the combined treatment of chemotherapy and USP7 inhibitor.

It is also worth noting that genetic deletion of *Usp7 in vivo* displays a stronger effect of tumor suppression than the treatment of USP7 inhibitor in PDOs. One likely explanation is the efficacy of the USP7 inhibitor P22077. Although several second generation USP7 inhibitors have been described with increased selectivity and potency (42–45), there is limited data on the use of these inhibitors *in vivo*. Given that Usp7 regulates both p53 and β-catenin, and potentially other substrates, it will be worth exploring if any of these (and other) new USP7 inhibitors target specifically the interaction between USP7 and β-catenin to increase the drug selectivity and efficacy.

In conclusion, our data provide robust *in vivo* evidence of USP7 as a tumor-specific therapeutic target in *APC*-mutated CRCs. Our finding is inconsistent with the previous suggestion of USP7 as a potent negative WNT regulator (31), at least not in the intestine. However, it should be noted that the current study focuses mainly on the intestinal tract. We cannot exclude the possibility that USP7 may play a different role in other systems such as osteoblast and adipocyte differentiation, as previously suggested (31). Further investigation will be needed to fully characterize the functional roles of USP7 in different tissue systems.

## EXPERIMENTAL PROCEDURES

### Antibodies and other reagents

β-catenin (610154, BD), caspase-3 (9661L, Cell Signaling), Cyclin D1 (2978S, Cell Signaling), Lysozyme (A0099, Dako), Sox9 (AB5335, Millipore), Olfm4 (39141S, Cell Signaling), USP7 (Bethyl laboratories) and Keratin20 (13063S, Cell Signaling) were used in immunohistochemistry analysis. DUB inhibitor VI, P22077 (Calbiochem) was resuspended in DMSO at 10 mM (for organoid experiments) or 15 mg/ml (for mice experiments).

### Real-time quantitative RT–PCR

RNA was extracted according to the manufacturer’s instructions (Qiagen RNAeasy). cDNA was prepared using Maxima first strand cDNA synthesis kit with dsDNase (#1672, Thermo Scientific). Quantitative PCR detection was performed using iTaq SYBR Green Supermix (#172-5121, Bio Rad) using specific primers to: *hβ-ACTIN* F: 5’ TTCTACAATGAGCTGCGTGTG 3’ R: 5’ GGGGTGTTGAAGGTCTCAAA 3’; *hCCND1*: F: 5’ CTCCGCCTCTGGCATTTTGG 3’ R: 5’ TCTCCTTGCAGCTGCTTAG 3’; *hAXIN2*: F: 5’ AGTGTGAGGTCCACGGAAAC 3’ R: 5’’
sCTTCACACTGCGATGCATTT 3’; *hASCL2*: F: 5’ GACCTGCGTACCTTGCTTTG 3’ R: 5’ CGCGCGATCACATTCTGTAA 3’; *hLGR5*: F: 5’ ACTGCATCCTAAACTGCCCT 3’ R: 5’ TGTCCAGACGTAGGTTTGCT 3’; *mUsp7*: F: 5’ TGCTGAATCTGACTCCACGT 3’ R: 5’ CCCAGTCGTTTTCCTTGTGG 3’; *mAxin2*: F:5’ TCCAGAGAGAGATGCATCGC 3’ R: 5’ AGCCGCTCCTCCAGACTATG 3’; *mAscl2*: F: 5’ AATGCAAGCTTGATGGACGG 3’ R: 5’ GGAAGCCCAAGTTTACCAGC 3’; *mLgr5*: F: 5’ CATCAGGTCAATACCGGAGC 3’ R: 5’ TAATGTGCGAGGCACCATTC 3’; *mHrpt1*: F: 5’TCATGAAGGAGATGGGAGGC 3’; *mOlfm4*: F: 5’ GACAGAGTGGAGCGCTTAGAG 3’ R: 5’ GCATCTCCTTCACTTCCAGC 3’; *mSox9*: F: 5’ CTGGAGGCTGCTGAACGAGAG 3’ R: 5’ CGGCGGACCCTGAGATTGC 3’. After cDNA amplification (40 cycles), samples were normalized to β-actin (human) or *Hrpt1* (mouse), and data were expressed as mean ± SD.

### Immunohistochemistry and Edu staining

For analysis of small intestine and colon by immunohistochemistry and Edu staining, tissues were fixed in 10% formalin and embedded in paraffin. For small intestinal tissues, same proximal part of the small intestine from all genotypes was used throughout the study. Immunohistochemistry was performed as described (30). The buffer used for antigen retrieval was citrate (Sox9, Olfm4, Keratin20 and Caspase-3). Edu staining was performed following manufacturer’s instructions (C10338, Invitrogen). Mouse adenomas were graded by a pathologist by analyzing H&E stained sections as previously described (46). Colitis and enteritis were graded by analysis of H&E sections by a pathologist as described in Fig. 3g, Fig. S3g, Fig. 4g and Fig. S4i). The following parameters were histopathologically assessed and scored for small intestine enteritis assessment: a) epithelial injury (including villous atrophy in small intestine, 0-3), b) lamina propria inflammation (0-3), c) area (% section) affected (0-3) and d) markers of severe inflammation (including crypt abscesses, submucosal inflammation and/or ulceration, 0-3). The total possible score is thus out of 12.

### RNAScope in situ hybridization

In situ hybridization (ISH) for Lgr5 was performed using the RNAScope FFPE assay kit (Advanced Cell Diagnostics) according to the manufacturer’s instructions. Briefly, 4 μm formalin-fixed, paraffin-embedded tissue sections were pre-treated with heat and protease digestion before hybridization with the target probe. Then, an HRP-based signal amplification system was hybridized to the target probes (Lgr5, 312171) before color development with 3,30-diaminobenzidine tetrahydrochloride (DAB). 20 crypts from 3 mice per group were used to quantitate the number of Olfm4+ cells.

### Mouse intestinal organoid culture

Organoids were established from freshly isolated wild-type (Control), Usp7 cKO, APC^fl/fl^ Control and APC^fl/fl^ Usp7 cKO small intestines. Tissues were incubated in cold PBS containing 2 mM EDTA for isolating epithelial crypts and culture as previously described (47) except that Matrigel was replaced with Cultrex BME, Type 2 RGF PathClear (Amsbio 3533-010-02). In brief, the organoid basal media contains EGF (Invitrogen PMG8043), Noggin and R-spondin (ENR) (5%). For APC^fl/fl^ Control and APC^fl/fl^ Usp7 cKO organoids, R-spondin was withdrawn from the media. The Rho kinase inhibitor Y-27632 (Sigma) was added to the culture when trypsinized.

### Human material and patient-derived organoid culture

Patient-derived organoids (PDOs) were derived from human CRC tissues that have been harvested during surgeries at the Royal Marsden Hospital (PDO1-2, REC reference 14/LO/1812), Guy’s and St. Thomas’ NHS Foundation Trust (PDO3, REC reference 12-EE-0493 and 18-EE-0025) and University College London Hospital (PDO4-5, REC reference 15/YH/0311) in accordance with ethical approval. Written informed consent was obtained. Intestinal samples were obtained from patients with colorectal cancer. Crypts were isolated from human intestinal tissue by incubating for 1 hour with chelation buffer (5.6 mM Na_2_HPO_4_, 8 mM KH_2_PO_4_, 96 mM NaCl, 1.6 mM KCl, 44 mM sucrose, 54.8 mM D-sorbitol, 0.5 M EDTA and 1M DTT at 4°C, and plated in drops of BME (47). After polymerization culture media was added. Human intestinal organoid media contains advanced DMEM/F12 medium (Invitrogen) including B27 (Invitrogen), nicotinamide (Sigma-Aldrich), N-acetylcysteine (Sigma-Aldrich), EGF (Invitrogen PMG8043), TGF-ß type I receptor inhibitor A83-01 (Tocris), P38 inhibitor SB202190 (Sigma-Aldrich), gastrin I (Sigma-Aldrich), WNT3a conditioned media (50% produced using stably transfected L cells), Noggin and R-spondin conditioned media. PDO1 and PDO2 were a kindly provided by Nicola Valeri’s Lab. More details from these two patients can be found in their publication (48).

### Organoid colony formation assay

Organoids were trypsinized and counted. 2,000 single cells were seeded in BME per 48 wells and placed in a 37°C incubator to polymerize for 20 min. 250 μl of complete Organoid Growth media plus Y-27632 was then added and cultured for 6 days for mouse organoids and 10 days for human organoids. Number of spheres formed in each well was counted as plating efficiency. Experiments were performed in triplicate.

### Cell Titer Glo

Organoids were trypsinized using TrypLE and filtered with a 20 μm cell strainer. 2000 single cells were seeded in BME per 48 well and placed in a 37°C incubator to polymerize for 20 minutes. 300 μl of complete Organoid Growth media was added for 10 days. The media was supplemented with 10 μM Y-27632 for 2 days after plating. Cell Titer Glo Luminiscent Cell viability assay (G7572, Promega) was used to assess viability of organoids. Experiments were performed at least three times with three triplicates each.

### Animal procedures

All animal regulated procedures were carried out according to Project License constraints (PEF3478B3 and 70/8560) and Home Office guidelines and regulations. In accordance with the 3Rs, the smallest sample size was chosen that could show a significant difference. Usp7^fl/fl^ mice were obtained from (32). Animals of both sexes at age 6–7 weeks on C57/BL6J background were used for the different experimental conditions and harvested as indicated.

Tamoxifen was injected intraperitoneally for 3 consecutive days (1.5 mg/10 g of mouse weight) from a 20 mg/ml stock solution. For experiments involving APC^min^ mice, animals were injected once a week with the same dose of tamoxifen after the first week of injections to ensure proper Usp7 deletion. For experiments with Villin^CreERT2^; Usp7^fl/fl^ mice that were let to age up to 1.5 years, one injection of the same dose of tamoxifen was injected once a month to ensure proper Usp7 deletion. For proliferation analysis, 5-ethynyl-2’-deoxyuridine (EdU) (Life Technologies) was injected intraperitoneally (0.3 mg/10 g of mouse weight) from a 10 mg/ml stock solution and mice were culled 2h after the injection.

### Xenograft experiments

PDO3 and PDO5 human organoids were subcutaneously injected into both flanks of 6-8-week-old NSG mice. The PDO-xenografts were allowed to grow 2-3 mm before randomizing the mice into a control group (DMSO) or P22077 (30 mg/kg) by intraperitoneal injection every day for 20 days. At the end of the experiments, all mice were culled. Tumors were collected, weighted, and photographed. All mice were housed in a pathogen-free environment and handled in strict accordance to institutional protocol.

### Quantification and statistical analysis

Statistical analyses were performed using GraphPad Prism 8 software. Statistical details and sample numbers are specified in the figure legends. For parametric data, statistical significance was determined using student’s unpaired, two-tailed t-test. For survival experiments, Log-rank (Mantel–Cox) test was used. P values are represented as *P < 0.05; **P < 0.01; ***P < 0.001.

## Acknowledgements

We thank the Francis Crick Institute’s Biological Research Facilities and Experimental Histopathology for managing mouse colonies and tissue processing. This work was supported by the Francis Crick Institute, which receives its core funding from Cancer Research UK (FC001105), the UK Medical Research Council (FC001105), and the Wellcome Trust (FC001105). Work in the V.S.W.L laboratory was also supported by the European Union’s Horizon 2020 research and innovation programme (668294). For the purpose of Open Access, the author has applied a CC BY public copyright license to any Author Accepted Manuscript version arising from this submission.

## Author contributions

L.N. designed the study, conducted the experiments, analyzed the data, and wrote the manuscript. A.K. and A.B. performed and analyzed the mouse experiments. G.V., D.R., F.C. and N.V. harvested human CRC tissues and generated PDOs. A.R. provided technical support to PDO characterization. V.S.W.L. designed the study and wrote the manuscript.

## Figure legends

**Figure S1.**
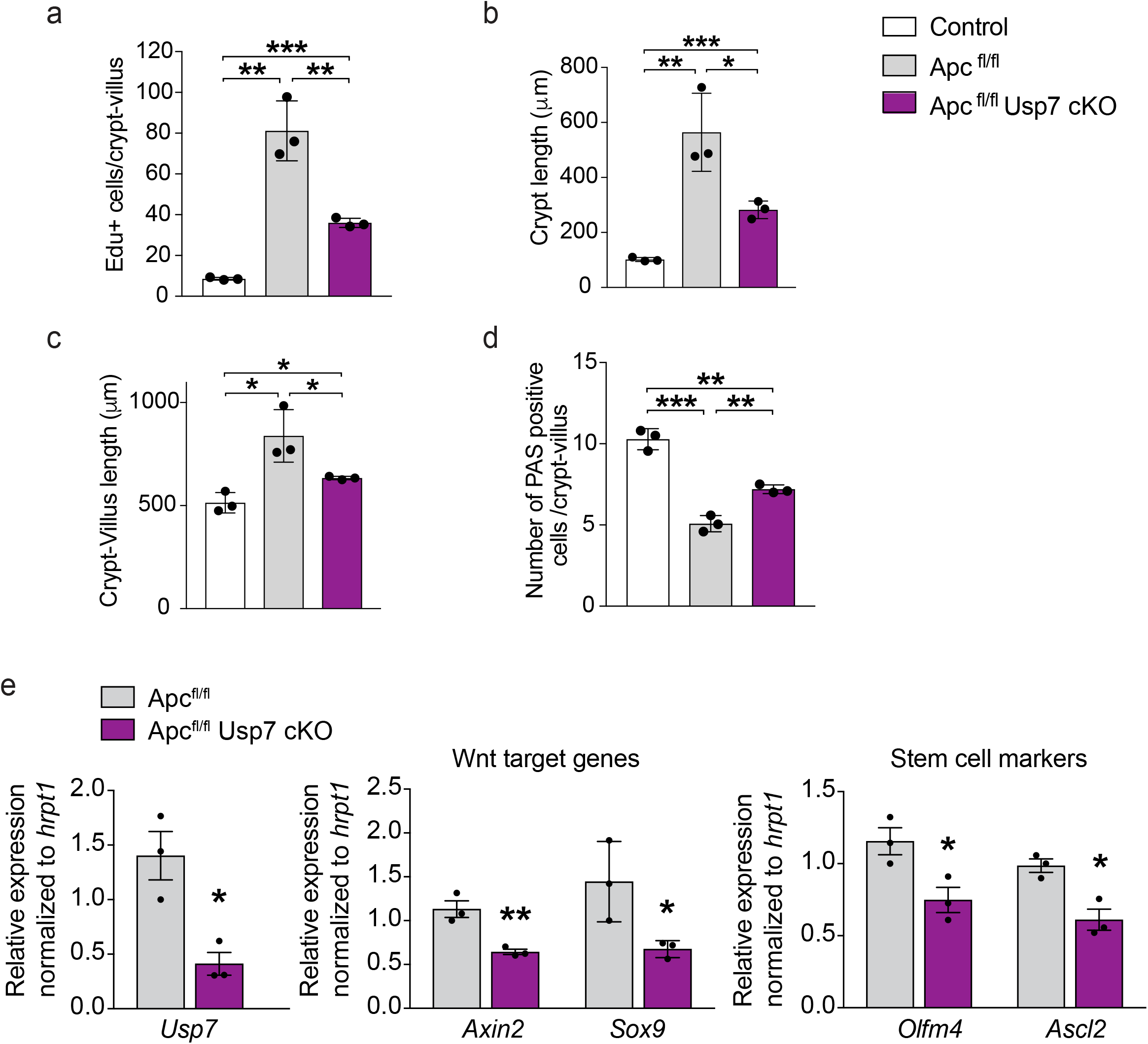
Usp7 inhibition decreases WNT signaling and proliferation in organoids derived from APC^fl/fl^ mice. (**a**) Quantitation of Edu positive cells from (Fig.1). Each dot represents the average of at least 10 crypts per animal. (**b**-**c**) Quantitation of crypt length (μm) (b) and crypt-villus length (μm) (c) from (Fig.1). Each dot represents the average of at least 10 crypts per animal. Data are mean ± standard error. n=3 per group. (**d**) PAS staining (indicative of Goblet cells) positive cells from (Fig.1). Each dot represents the average of at least 20 crypts per animal. (**e-f**) mRNA expression of the indicated genes was analyzed by qRT-PCR in small intestinal organoids from (Fig. 1v). Data are presented as fold change normalized to *hrpt1* (n=3 per condition). Error bars represent ± standard error. n=3 per group. P-values were determined using the unpaired two-sided t-test (*P<0.05; **P<0.01; ***P<0.001).

**Figure S2.**
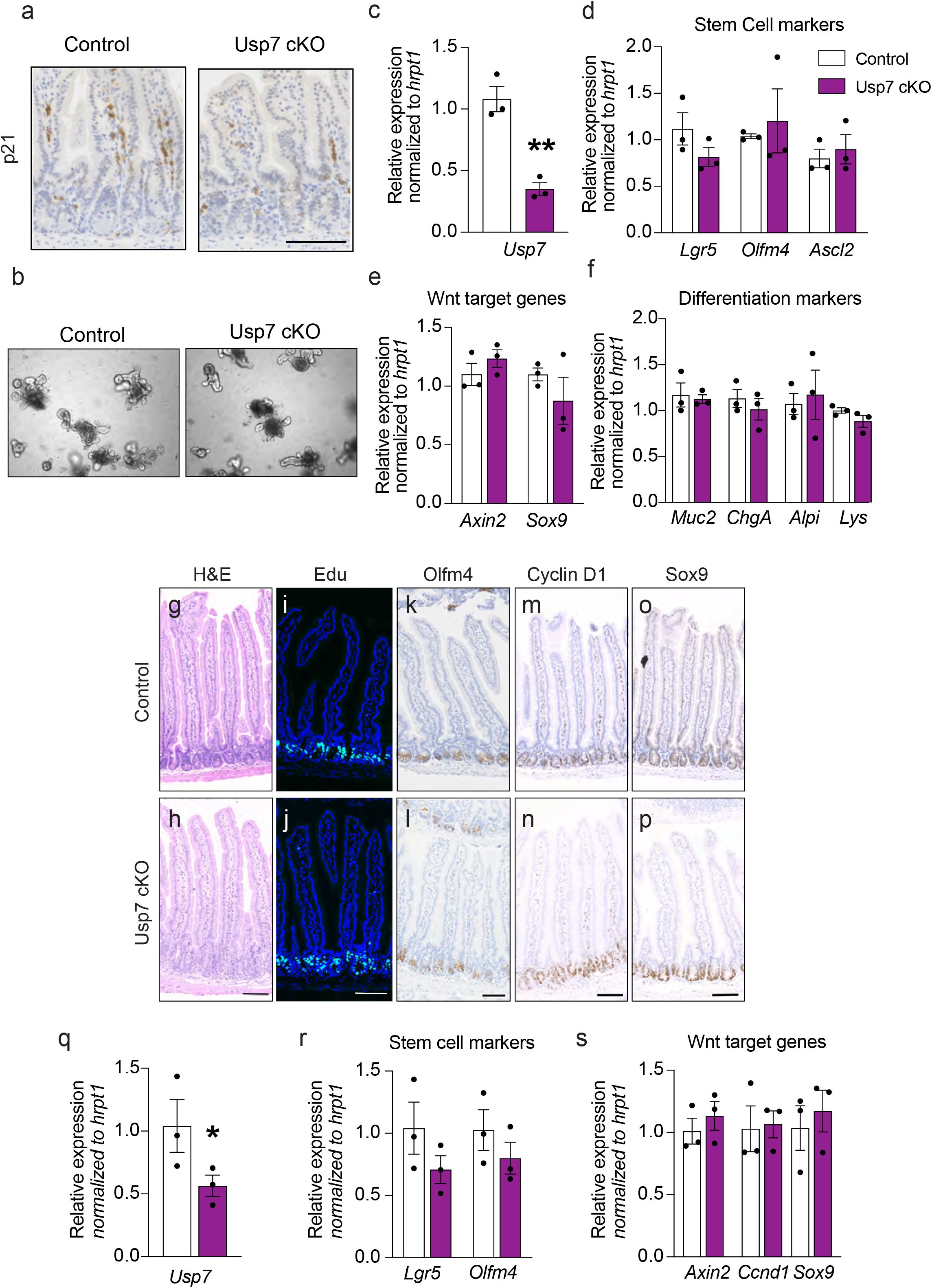
Loss of Usp7 in wild-type APC mice maintains intestinal homeostasis. **(a)** Representative images of p21 staining in Villin^CreERT2^ (Control) and Villin^CreERT2^; Usp7^fl/fl^ (Usp7 cKO) intestine. Scale bar, 100 μm. **(b)** Representative images of organoids derived from control and Usp7 cKO mice at 7 dpi. Scale bar, 100 μm. (**c-f**) mRNA expression of the indicated genes was analyzed by qRT-PCR from organoids isolated from Control and Usp7 cKO mice. Data are presented as fold change normalized to *β-actin* control in triplicate (n=3 per condition). (**g**-**p**) Representative images of H&E staining (g-h), Edu (i-j), Olfm4 (k-l), Cyclin D1 (m-n), Sox9 (o-p) from Control and Usp7 cKO intestine 1.5 years post-tamoxifen induction. Scale bars, 50 μm. n=3 per group. (**q**-**s**) mRNA expression of the indicated genes was analyzed by qRT-PCR in crypts from 1.5 years old Control and Usp7 cKO mice. Data are presented as fold change normalized to *hrpt1* (n=3 per condition). Error bars represent ± standard error. n=3 per group. P-values were determined using the unpaired two-sided t-test (*P<0.05; **P<0.01; ***P<0.001).

**Figure S3.**
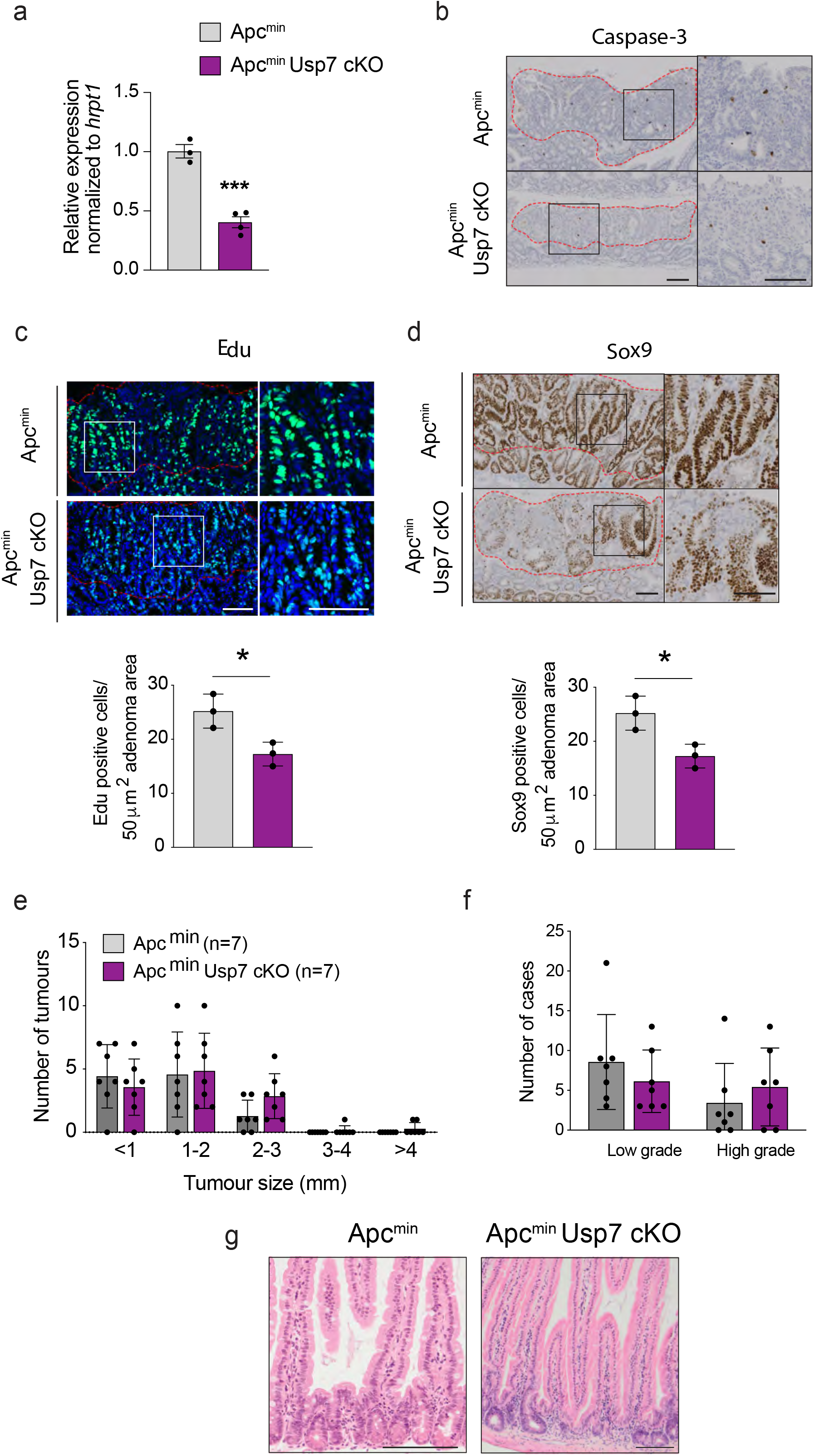
Analysis of tumors derived from Apc^min^ and Apc^min^ Usp7 cKO mice. (**a**) mRNA expression of the Usp7 was analyzed by qRT-PCR in crypts from Apc^min^ Control and Apc^min^ Usp7 cKO mice. Data are presented as fold change normalized to *hrpt1* (n=3 per condition). Error bars represent ± standard error. P-values were determined using the unpaired two-sided t-test (***P<0.001). (**b**-**d**) Representative Caspase-3 (b), Edu (c) and Sox9 (d) stainings of tumors derived from the indicated genotypes 120 days after Usp7 loss. Graphs in (c) and (d) indicate the number of Edu (c) or Sox9 (d) positive cells in 50 μm^2^ of adenoma area. n=3 per group. Scale bar, 100 μm. (**e-f**) Quantitation of the total number of tumors (e) and tumor grades (f) developed in the indicated mice at around 200 days after the first tamoxifen induction. Data are mean ± standard error. Apc^min^ control, n=7; Apc^min^ Usp7 cKO, n=7. (g) Representative H&E staining from Apc^min^ control and Apc^min^ Usp7 cKO mice showing enteritis present in the Apc^het^ Usp7 cKO mice. Scale bar, 100 μm.

**Figure S4.**
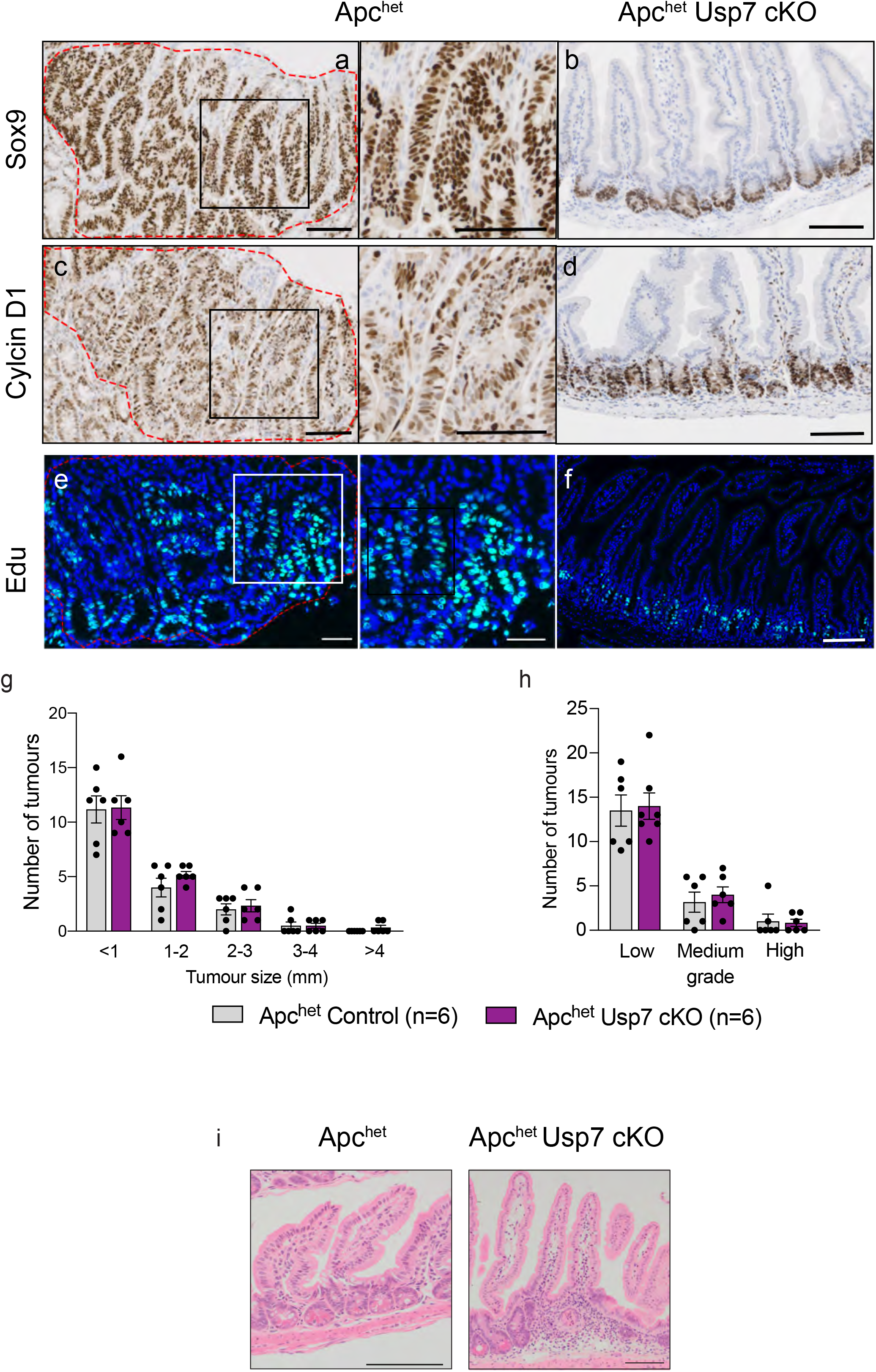
Analysis of tumors derived from Apc^het^ and Apc^het^ Usp7 cKO mice. (**a-f**) Immunostaining of Apc^het^ (Control) tumors (a, c, e) and Apc^het^ Usp7 cKO (b, d, f) untransformed intestine 97 dpi using the indicated antibodies (n=3 per group). Scale bars, 50 μm. (**g-h**) Quantitation of the total number of tumors (g) and tumor grades (h) developed in the indicated mice at around 200 days after the first tamoxifen induction. Data are mean ± standard error. Apc^het^ control, n=6; Apc^het^ Usp7 cKO, n=6. (i) Representative H&E staining from Apc^het^ control and Apc^het^ Usp7 cKO mice showing enteritis present in the Apc^het^ Usp7 cKO mice. Scale bar, 100 μm.

**Figure S5.**
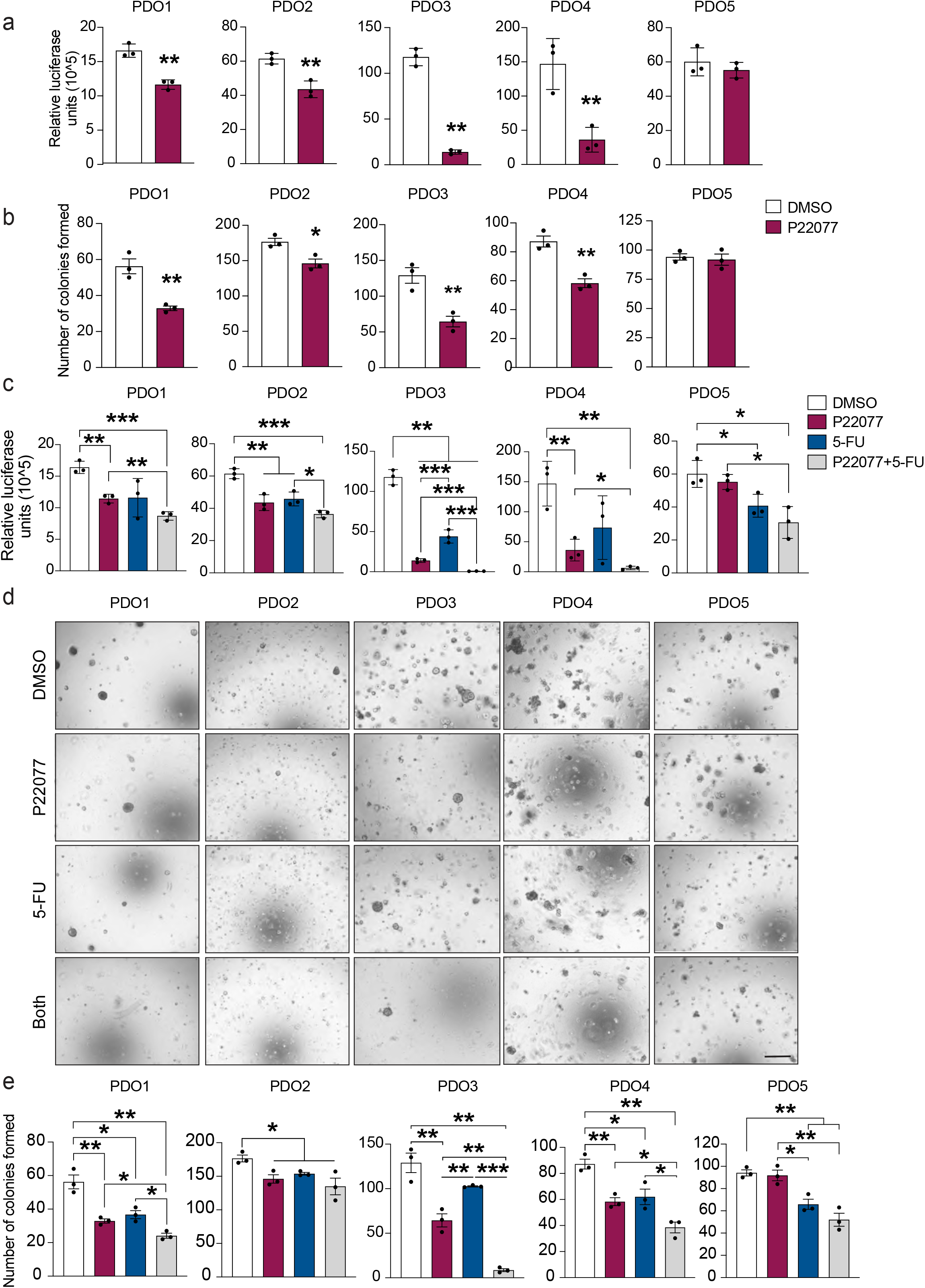
Inhibition of USP7 in human CRC patient-derived organoids suppresses cell growth and sensitizes 5-FU treatment. (**a**) Cell Titer Glo luciferase assay of the indicated PDOs treated with P22077 or vehicle (DMSO). Error bars represent ± standard error from three independent experiments. (**b**) Quantitation of the number of colonies formed in the indicated PDOs treated with DMSO or P22077 after 10 days. Data are mean ± standard error. At least n=3 per group. (**c**) Cell Titer Glo luciferase assay of the indicated organoids treated with DMSO, P22077, 5-FU or both for 10 days. Error bars represent ± standard error from three independent experiments. (**d**) Representative images of colony formation assay of organoids from the organoids in (c). Scale bar, 1000 μm. (**e**) Quantitation of the number of colonies formed in the indicated PDOs in (d) treated with DMSO, P22077, 5-FU of both after 10 days. Data are mean ± standard error. At least n=3 per group. P-values were determined using the unpaired two-sided t-test (*P<0.05; **P<0.01; ***P<0.001).

**Figure S6.**
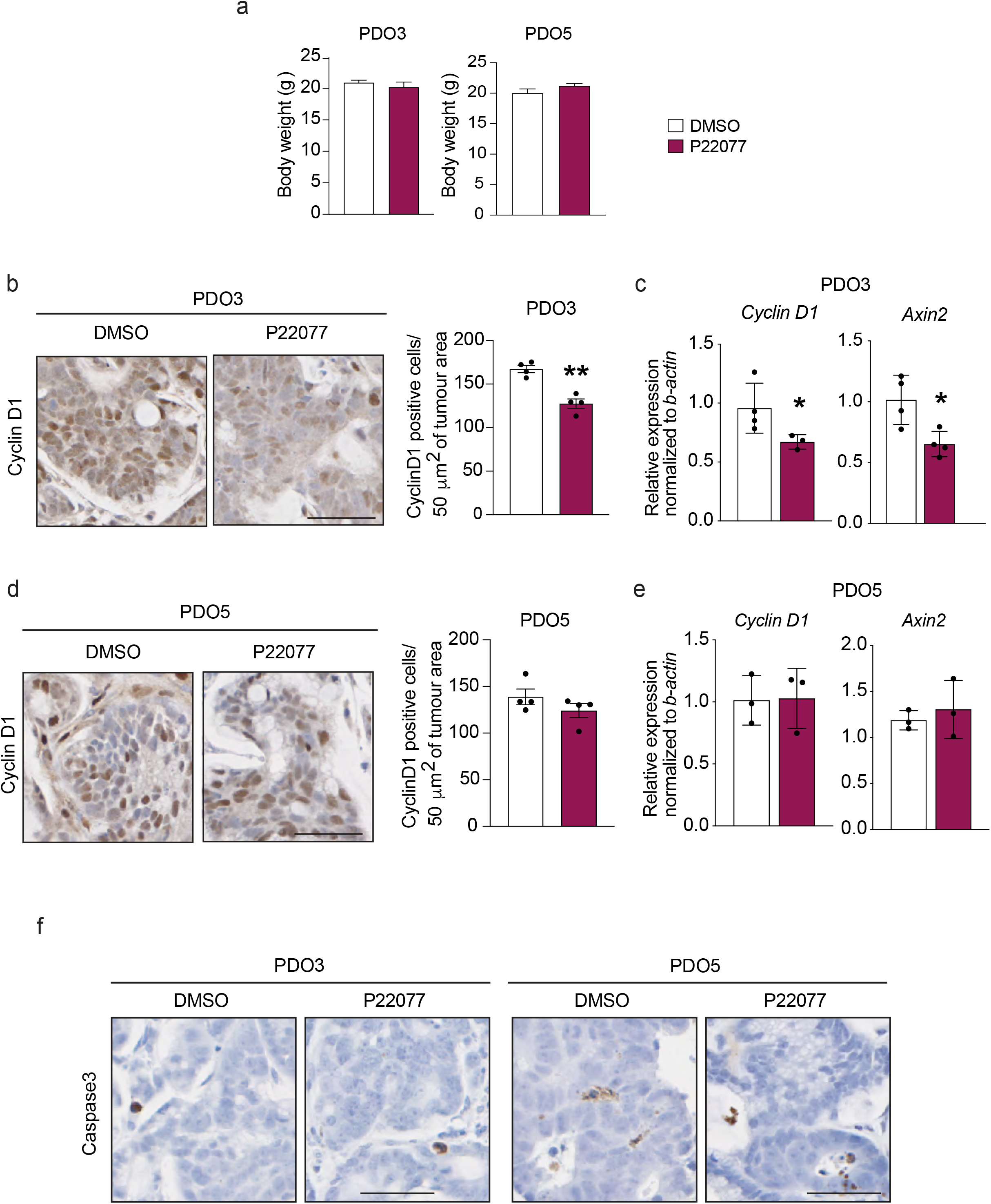
Inactivation of USP7 inhibits WNT signaling in APC-truncated PDO-derived xenografts *in vivo*. (**a**) Measurement of body weight of mice transplanted with PDO3 or PDO5 and treated with DMSO or P22077 inhibitor (30 mg/kg) at the end of the experiment (20 days). (**b, d**) Representative images of Cyclin D1 stainings from DMSO- or P22077-treated PDO3 (b) or PDP5-derived (d) tumors. Scale bar, 50 μm. Left panel: Quantitation of Cyclin D1 positive cells shown on the right panel. Each dot represents the average of at least 6 different areas of 50 μm^2^ per tumor. Data are mean ± standard error. n= 4 per group. P-values were determined using the unpaired two-sided t-test (**P<0.01). (**c, e**) mRNA expression of the indicated genes was analyzed by qRT-PCR in tumors derived from PDO3 (c) and PDO5 (e). Data are presented as fold change normalized to *β-actin*. Error bars represent ± standard error. At least n=3 per group. P-values were determined using the unpaired two-sided t-test (*P<0.05). (**f**) Immunohistochemical analysis of PDO3- and PDO5-derived xenografts treated with DMSO or P22077 and stained with cleaved Caspase 3. Scale bar, 50 μm.

